# GluK2 Q/R editing regulates kainate receptor signalling to modulate AMPA receptor synaptic expression and plasticity

**DOI:** 10.1101/2022.10.31.514576

**Authors:** Jithin D. Nair, Kevin A. Wilkinson, Busra P. Yucel, Christophe Mulle, Bryce Vissel, Jack Mellor, Jeremy M. Henley

## Abstract

Q/R editing of the kainate receptor (KAR) subunit GluK2 radically alters properties of recombinant KARs, but the effects *in vivo* remain largely unexplored. We compared GluK2 editing-deficient mice that express ∼95% unedited GluK2(Q) to wild-type counterparts that express ∼85% edited GluK2(R). At mossy fibre-CA3 (MF-CA3) synapses GluK2(Q) mice displayed enhanced postsynaptic KAR function and increased KAR-mediated presynaptic facilitation, demonstrating heightened ionotropic function. Conversely, there was reduced metabotropic KAR function, assessed by KAR-mediated afterhyperpolarization currents, in GluK2(Q) mice. GluK2(Q) mice had fewer GluA1- and GluA3-containing AMPA receptors (AMPARs) and reduced postsynaptic AMPAR currents at both MF-CA3 and CA1-Schaffer collateral synapses. Moreover, long-term potentiation of AMPAR-mediated transmission at CA1-Schaffer collateral synapses was reduced in GluK2(Q) mice. These findings suggest that GluK2 Q/R editing influences ionotropic/metabotropic balance of KAR signalling to regulate synaptic expression of AMPARs and plasticity.

## Introduction

Kainate receptors (KARs) are glutamate-gated cation channels assembled from tetrameric combinations of the subunits GluK1-GluK5. Depending on the synapse and neuron type, KARs are present at both pre- and postsynaptic sites throughout the brain ^1^. Despite their close homology to AMPA- and NMDA-type glutamate receptors, postsynaptic KARs mediate only a minor fraction of the ionotropic synaptic response to glutamate but they are critically important for synaptic integration and regulation of neural circuits ^2–4^. Presynaptic KARs also contribute to neuronal network function by regulating neurotransmitter release probability at both excitatory and inhibitory synapses ^5–11^.

KARs have been particularly well studied at hippocampal glutamatergic MF-CA3 synapses ^12, 13^, where the GluK2 subunit is an integral component of both pre- and postsynaptic KARs ^6–8, 14–16^. Indeed, ionotropic postsynaptic KARs containing GluK2 were first discovered at MF-CA3 synapses ^12,13,17^, and GluK2-containing KARs were shown to contribute to short-term plasticity of presynaptic release probability over timescales ranging from 10ms to 20s ^18,19^.

In addition to ionotropic actions, KARs also initiate G protein-coupled metabotropic signalling ^20–24^. Pharmacological activation of KARs by exogenous agonists regulates presynaptic release of both GABA and glutamate through this metabotropic pathway ^20,25^. Metabotropic signalling through postsynaptic KARs has been demonstrated at Schaffer collateral CA1 synapses ^26^ and at MF-CA3 synapses ^15,27^. Synaptic activation of postsynaptic KARs inhibits the slow after hyperpolarization (I_sAHP_), a long-lasting voltage-independent and Ca^2+^-dependent K^+^ current produced following short bursts of action potentials ^28^. KAR-mediated inhibition of I_sAHP_ occurs in multiple neuronal types via a G_i/o_ G protein and PKC-dependent pathway and, in CA3 cells, is absent in GluK2 knockout mice, suggesting a crucial role for this subunit in this form of metabotropic signalling ^15,26,27,29^.

KAR surface expression is activity-dependently and bidirectionally regulated ^24,30–34^. Furthermore, activation of KARs can also up- or down-regulate AMPAR surface expression to mediate plasticity ^24,35^. Specifically, transient activation of GluK2-containing KARs in cultured hippocampal neurons increases AMPAR surface expression, and in hippocampal slices GluK2-containing KARs can induce AMPAR long-term potentiation (KAR-LTP_AMPAR_) at Schaffer collateral-CA1 synapses via a pertussis toxin-sensitive metabotropic signalling pathway ^24^. In contrast, sustained activation of KARs reduces surface expression of AMPARs in cultured hippocampal neurons and induces AMPAR long-term depression in CA1 neurons in hippocampal slices (KAR-LTD_AMPAR_), an effect that is lost in the absence of GluK2 ^35^. These results highlight the importance of GluK2-containing KARs as modulators of AMPAR-mediated synaptic transmission.

The nuclear enzyme ADAR2 Q/R edits GluK2 pre-mRNA, resulting in a genetically encoded glutamine (Q) in the channel pore region being replaced by an arginine (R) ^36,37^. In recombinant systems KARs containing GluK2(R) subunits display ER retention and reduced traffic to the cell surface compared to those assembled with the unedited GluK2(Q) ^38,39^. Furthermore, the edited GluK2(R)-containing KARs that do reach the surface do not gate Ca^2+^ and have a channel conductance <1% of GluK2(Q) ^40^.

The dynamic regulation of GluK2 Q/R editing underpins KAR homeostatic plasticity whereby chronic suppression of network activity decreases ADAR2 levels. This, in turn, reduces GluK2 editing resulting in enhanced KAR surface expression. Reciprocally, chronic enhancement of network activity promotes GluK2 Q/R editing and reduces KAR surface expression ^32–34^.

We used electrophysiology and biochemistry experiments approaches to investigate how GluK2 Q/R editing alters KAR signalling and function in intact neuronal circuits and whether these changes, in turn, regulate AMPAR function. GluK2 editing-deficient mice contain a deletion in the intronic editing complementary sequence (ECS) of the *grik2* gene that directs ADAR2-mediated codon substitution in the GluK2 pre-mRNA (Figure 1A). This results in >95% of GluK2-KARs in adult mice containing unedited GluK2(Q), whereas KARs in WT mice contain <15% GluK2(Q) ^41^. GluK2(Q) mice are viable and, as expected, surface expressed KARs in cultures from these mice display the inwardly rectifying current/voltage (I-V) relationship characteristic of GluK2(Q)-containing KARs expressed in recombinant systems ^42,43^. The only previous study using these GluK2(Q) mice detected no differences in GluK2 mRNA levels, no alterations in editing of the other GluK2 editing sites (I/V and Y/C) and no change in Q/R editing of the GluA2 AMPAR subunit ^41^. However, the GluK2(Q) mice were reported to have increased susceptibility to kainate-induced seizures and display a form of NMDAR-independent LTP at the medial perforant-DG synapses that is not present in WT mice ^41^.

**Figure 1:**
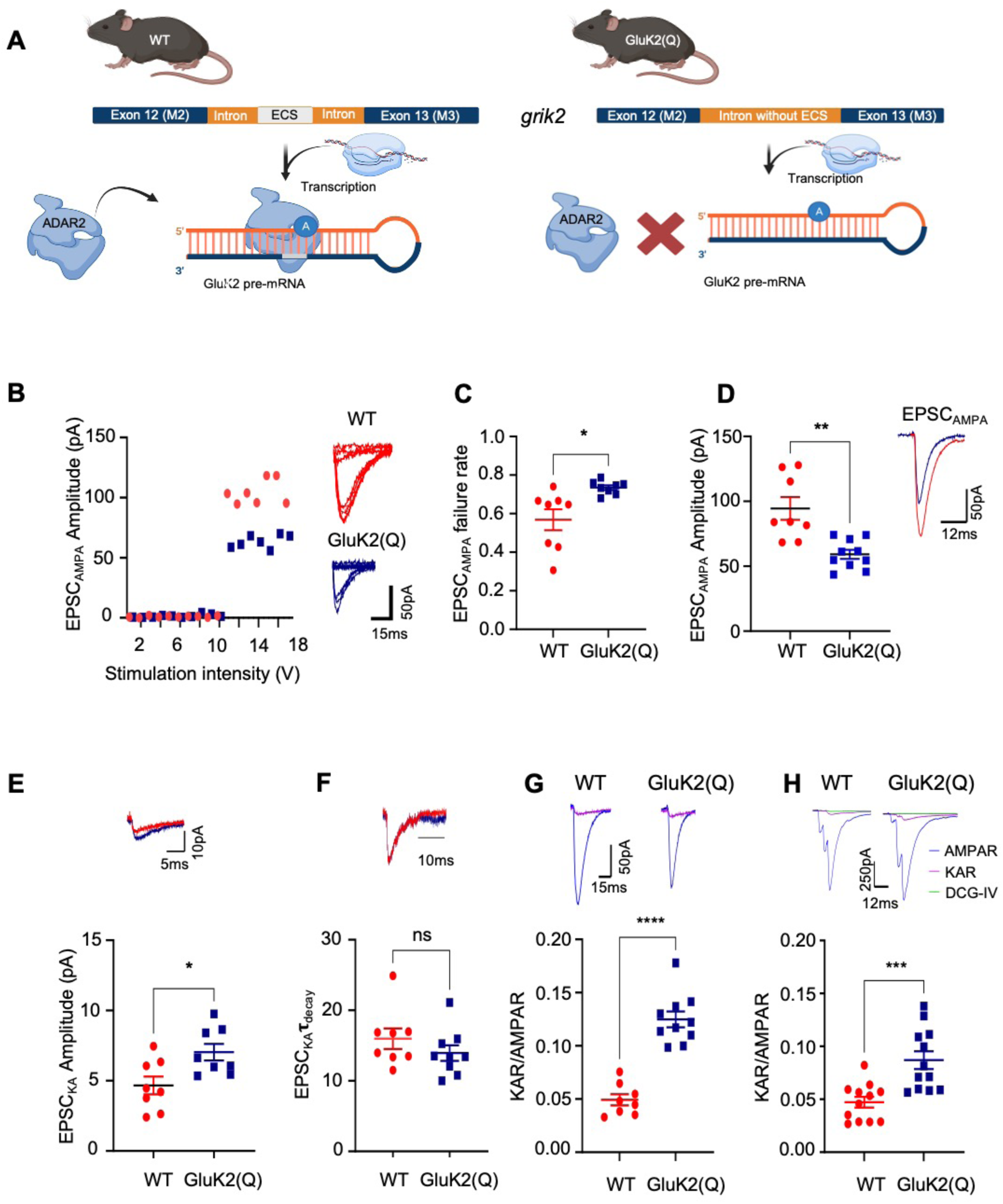
Enhanced postsynaptic KAR and reduced AMPAR currents at MF-CA3 synapses in GluK2(Q) mice. **(A)** The intronic region between exon 12 (M2) and 13 (M3) in the *grik2* gene contains an editing complementary site (ECS), located ∼1900nt downstream of exon 12. In the WT mice, this region is intact. However, in homozygous GluK2(Q) mice, a 600bp region is deleted from the ECS site. This prevents ADAR2 binding and subsequent editing of GluK2 pre-mRNA at the Q/R site, leading to the translation of >95% unedited GluK2(Q) subunits. **(B)** Representative data from a minimal stimulation experiment showing stimulation intensity threshold for evoking responses from excitation of a single axon fibre (left). Eight superimposed consecutive traces showing synaptic successes and failures for minimal stimulation for WT (top right) and GluK2(Q) mice (bottom right). **(C)** Quantification of probability of failures for AMPAR responses out of 150 stimulations in WT and GluK2(Q) mice. N=5, n=8 cells for WT and N=5, n=10 cells for GluK2(Q) mice; ns p>0.05, *p<0.05, **p<0.01; unpaired t-test with Welch’s correction. **(D)** Quantification of average amplitude of EPSC_AMPA_ (left). Representative trace showing EPSC_AMPA_ in WT and GluK2(Q) mice (right).N=5, n=8 cells for WT and N=5, n=10 cells for GluK2(Q) mice (bottom).; ns p>0.05, *p<0.05, **p<0.01; unpaired t-test with Welch’s correction. **(E)** Representative trace showing EPSC_KA_ in WT and GluK2(Q) mice (top). Quantification of average amplitude of EPSC_KA_ (bottom). N=5, n=8 cells for WT and N=5, n=10 cells for GluK2(Q) mice (bottom); ns p>0.05, *p<0.05, **p<0.01; unpaired t-test with Welch’s correction. **(F)** Representative trace showing !_decay_ for KAR currents (scaled) in WT vs GluK2(Q) (top) Quantification of !_decay_ for KAR currents in WT vs GluK2(Q) mice. N=5, n=8 cells for WT and N=5, n=10 cells for GluK2(Q) mice (bottom); ns p>0.05, *p<0.05, **p<0.01; unpaired t-test with Welch’s correction. **(G)** Representative traces showing postsynaptic AMPAR and KAR currents in WT and GluK2(Q) mice with minimal stimulation (top). Quantification of KAR/AMPAR current ratio in WT and GluK2(Q) mice (bottom) N=5, n=8 cells for WT and N=5, n=10 cells for GluK2(Q) mice, ns p>0.05, ****p<0.0001; unpaired t-test with Welch’s correction. **(H)** Representative traces showing postsynaptic AMPAR and KAR currents in WT and GluK2(Q) mice with burst stimulation at 167Hz (top). Quantification of KAR/AMPAR current ratio in WT and GluK2(Q) mice (bottom). N=5, n=12 cells, ns p>0.05, *p<0.05, **p<0.01; unpaired t-test with Welch’s correction.

Here, we show enhanced pre- and postsynaptic KAR ionotropic function in the GluK2(Q) mice compared to WT controls whereas metabotropic KAR-mediated inhibition of I_sAHP_ was reduced. Furthermore, in GluK2(Q) mice AMPAR-mediated transmission at hippocampal CA3 and CA1 synapses, synaptic levels of GluA1 and GluA3 were decreased and LTP at CA1 Schaffer collateral synapses was severely attenuated. We interpret these data to indicate that GluK2 editing is important for determining the mode of KAR signalling, and that this, in turn, plays key roles in the regulation of basal expression of synaptic AMPARs and AMPAR-mediated synaptic plasticity.

## Results

### Enhanced postsynaptic KAR and reduced AMPAR currents at MF-CA3 synapses in GluK2(Q) mice.

In recombinant systems KARs containing unedited GluK2(Q) have a higher conductance than edited GluK2(R) ^40,44^. Moreover, disrupting ADAR2-mediated GluK2 Q/R editing enhances the surface expression, single channel conductance, and Ca^2+^ permeability of postsynaptic KARs in cultured neurons ^33,40,45^. Therefore, we measured basal KAR-mediated postsynaptic responses at MF-CA3 synapses in WT and GluK2(Q) mice. Because both the glutamate receptor complement and presynaptic facilitation at MF-CA3 synapses are stable after the 2^nd^ postnatal week ^46,47^ we used P14-P21 mice and confirmed the stability of AMPAR and KAR expression within this age range.

To compare between GluK2-editing deficient and WT genotypes, we used minimal stimulation to isolate the synaptic response evoked by stimulation of a single presynaptic mossy fibre axon at 3Hz in acutely prepared hippocampal slices ^46^. Postsynaptic responses were recorded from CA3 pyramidal neurons in whole-cell voltage clamp configuration in the presence of picrotoxin (50µM) and D-APV (50µM) to block GABA_A_Rs and NMDARs, respectively. The stimulation intensity applied to presynaptic axons was increased in small increments until a postsynaptic response was observed (Figure 1B). Interestingly, the percentage of trials that evoked a synaptic response (success rate) was decreased in the GluK2(Q) mice (Figure 1C) (WT=43 ± 5%, GluK2(Q)=27 ± 1%; unpaired t-test, p=0.017), suggesting either a decrease in the number of release sites or reduced probability of glutamate release from the same number of sites. Furthermore, the average AMPAR-EPSC amplitude for successful trials was reduced in GluK2(Q) mice (Figure 1D) (WT=94.5 ± 8.8pA, GluK2(Q)=59.1 ± 3.5pA; unpaired t-test, p=0.0044).

The AMPAR antagonist GYKI53655 (40µM) was then applied to pharmacologically isolate KAR-mediated excitatory postsynaptic currents (EPSC_KA_) evoked by single presynaptic mossy fibre axons. Analysis of detectable synaptic responses revealed that EPSC_KA_ was increased in amplitude in GluK2(Q) mice (Figure 1E) (WT=4.66 ± 0.63pA, GluK2(Q)=7.31 ± 0.50pA; unpaired t-test, p=0.0057), but with no change in decay kinetics (Figure 1F) (!_decay,_ WT=16.0 ± 1.5ms, GluK2(Q)=13.9 ± 1.1ms; unpaired t-test, p=0.28), consistent with enhanced conductance of unedited GluK2(Q)-containing recombinant KARs ^40,44^.

We also measured the EPSC_KA_ - to EPSC_AMPA_ amplitude ratio for both minimal and larger EPSCs evoked by bursts of 3 stimuli at 167Hz given to mossy fiber axons that stimulate multiple axons. EPSC_AMPA_ was first collected in the presence of picrotoxin (50µM) and D-APV (50µM) to block GABA_A_Rs and NMDARs, respectively. Then, EPSC_AMPA_ was blocked by bath application of GYKI53655 (40µM) for 10 mins to isolate EPSC_KA_. The EPSC_KA_/EPSC_AMPA_ ratio increased in GluK2(Q) mice (Figure 1G, H) (minimal stimulation: WT=0.049 ± 0.005, GluK2(Q)=0.12 ± 0.007; unpaired t-test, p= 0.0001; 167Hz stimulation: WT=0.047 ± 0.005, GluK2(Q)= 0.086 ± 0.008; unpaired t-test, p=0.0008). Based on the minimal stimulation data for EPSC_KA_ and EPSC_AMPA_, this increase in KAR/AMPAR ratio likely results from an increase in EPSC_KA_, and a decrease in EPSC_AMPA_.

### Enhanced presynaptic facilitation in GluK2(Q) mice

Presynaptic KARs at MF-CA3 synapses are autoreceptors activated by released glutamate to facilitate the probability of vesicle release in response to subsequent action potentials ^8,48–50^. To investigate how GluK2 Q/R editing affects presynaptic KAR function at MF-CA3 synapses, we measured short-term facilitation of presynaptic release in the presence of picrotoxin (50µM) across a range of timescales. By measuring EPSC_AMPA_, we examined paired-pulse facilitation (PPF) at 50ms stimulation intervals and accumulation of frequency facilitation (FF) at 1s stimulation intervals in acute slices from WT and GluK2(Q) mice. Both PPF and FF were increased in GluK2(Q) mice (Figure 2A, B) (PPF: WT=3.3 ± 0.2, GluK2(Q)=5.3 ± 0.6; un-paired t-test, p=0.0067; FF: WT=4.5 ± 0.4, GluK2(Q)=6.6 ± 0.6; unpaired t-test, p=0.0114), consistent with enhanced presynaptic KAR function and/or decreased basal release probability.

**Figure 2:**
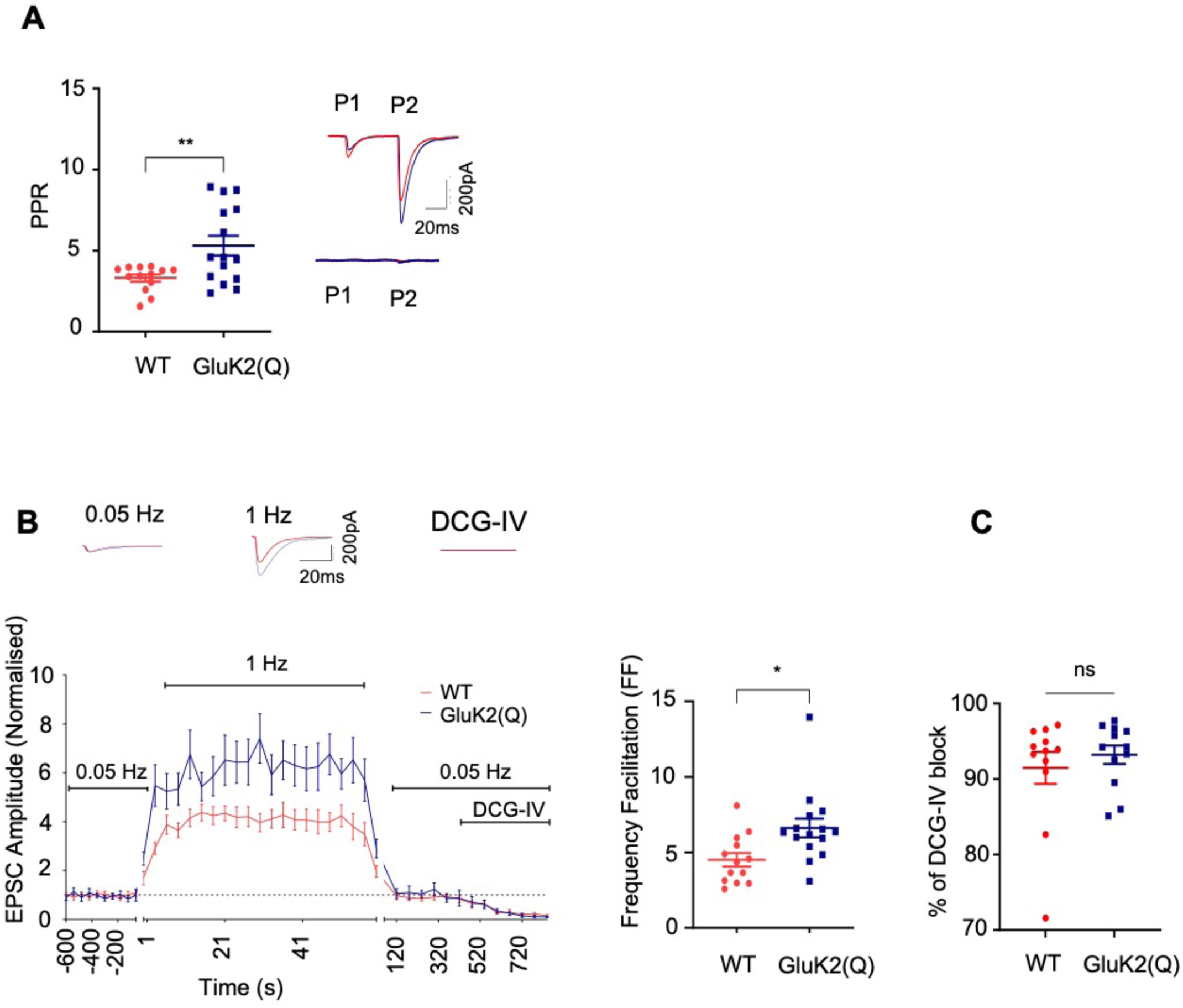
Enhanced presynaptic facilitation in GluK2(Q) mice. **(A)** Quantification of average paired-pulse ratio in WT and GluK2(Q) mice (left). Representative traces showing EPSCs in response to paired-pulse stimulation from both WT and GluK2(Q) mice before (top right) and after DCG-IV application (bottom right). WT, N=7, n=13 cells; GluK2(Q) mice, N=8, n=15 cells; ns p>0.05, *p<0.05, **p<0.01; unpaired t-test with Welch’s correction. **(B)** Representative trace showing frequency facilitation from WT and GluK2(Q) mice (top). Timeline of frequency facilitation experiments in WT and GluK2(Q) mice (bottom left). Quantification of frequency facilitation in WT and GluK2(Q) mice (bottom right). **(C)** Percentage of DCG-IV block in WT and GluK2(Q) mice. WT, N=7, n=13 cells; GluK2(Q) mice, N=8, n=15 cells; ns p>0.05, *p<0.05, **p<0.01; unpaired t-test with Welch’s correction.

In all experiments the purity of mossy fibre input was determined by addition of the group II metabotropic glutamate receptor agonist DCG-IV (2µM) ^51^ and recordings were excluded if there was <70% inhibition of synaptic responses. Furthermore, to exclude the possibility that differences in the purity of mossy fibre inputs contributed to the enhanced short-term facilitation in GluK2(Q) mice we analysed the degree of DCG-IV inhibition. No differences were observed (Figure 2C) (WT=91.5 ± 2.1%, GluK2(Q)=93.2 ± 1.2%; unpaired t-test, p=0.4879) and there was no correlation between degree of inhibition by DCG-IV and PPR (Supplementary Figure-S1A, B) (WT, r=0.277, R^2^=0.0769, p=0.383; GluK2(Q), r=-0.280, R^2^=0.0784, p=0.378; 95% confidence interval). There was also no correlation between EPSC amplitude and PPR (Supplementary Figure-S1C, D) (WT, r=0.127, R^2^=0.0163, p=0.692; GluK2(Q), r=0.0383, R^2^ = 0.00147, p=0.906; 95% confidence interval). EPSC initial amplitudes in response to the first stimulus were set to be similar between genotypes (WT=156.6 ± 14.4pA, GluK2(Q)=143.9 ± 17.8pA; unpaired t-test, p=0.5830).

Taken together with the reduced success rate observed in the minimal stimulation experiments (Figure 1C), these data suggest that basal release probability at MF-CA3 synapses is reduced in GluK2(Q) mice and that presynaptic facilitation is enhanced.

### Metabotropic KAR function is impaired in GluK2(Q) mice

Because the GluK2 Q/R editing site is within the channel pore region it is unsurprising that editing impacts on KAR ionotropic signalling but the effects of Q/R editing on metabotropic signalling have not been investigated. We therefore assessed if GluK2 Q/R editing alters metabotropic function by measuring KAR inhibition of the slow afterhyperpolarization current (I_sAHP_) in acute hippocampal slices ^15,26,27,52^. I_sAHP_ currents were evoked in whole-cell voltage clamped CA3 pyramidal cells by depolarising the membrane potential to 0mV from -50mV for 200ms in the presence of 50µM picrotoxin, 50µM D-APV, 40µM GYKI53655 and 1µM CGP55845 to inhibit GABA_A_Rs, NMDARs, AMPARs and GABA_B_Rs, respectively ^27^. Robust and stable I_sAHP_ currents were obtained in both WT and GluK2(Q) mice, with no difference in baseline amplitudes between genotypes (Figure 3A) (WT=67.0 ± 8.0pA, GluK2(Q)=63.1 ± 6.6pA; unpaired t-test, p=0.712). Activation of synaptic KARs by MF stimulation (10 stimuli at 25Hz every 20s for 10mins) ^27^ produced a consistent depression of I_sAHP_ in WT mice but the extent of depression was decreased in GluK2(Q) mice (Figure 3B) (WT=45.1 ± 3.7%, GluK2(Q)=26.7 ± 2.7%; unpaired t-test, p=0006). This data indicates that KAR metabotropic signalling is impaired in GluK2(Q) mice.

**Figure 3:**
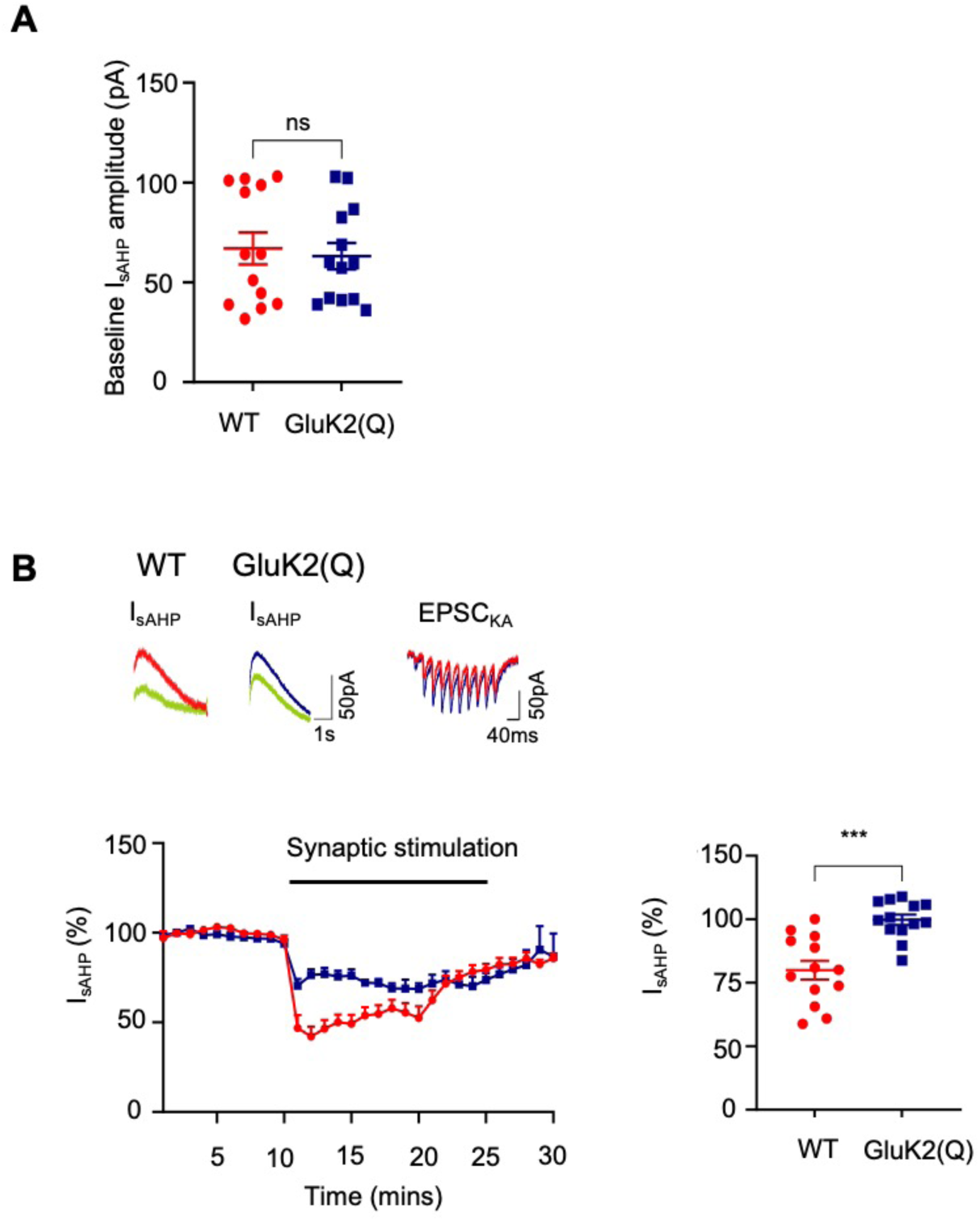
Impaired metabotropic KAR function in GluK2(Q) mice. **(A)** Average baseline amplitude of I_sAHP_ currents in WT and GluK2(Q) mice. **(B)** Sample traces from WT (top left) and GluK2(Q) mice (top middle) and EPSC_KA_ following synaptic stimulation (top right). Timeline showing inhibition of I_sAHP_ following synaptic KAR activation (bottom left). Quantification of percentage inhibition of I_sAHP_ following synaptic KAR activation in WT and GluK2(Q) mice (bottom right). N=4 animals, n=13 cells, ns p>0.05, **p<0.0001; un-paired t-test with Welch’s correction.

### Synaptic expression of KAR subunits is altered in GluK2(Q) mice

The reduced metabotropic signalling in GluK2(Q) mice could arise from disrupted molecular signalling and/or reduced expression of synaptic KARs. We therefore assessed total and synaptosomal expression of key KAR subunits in WT and GluK2(Q) mice by Western blotting. Total expression of the GluK1, GluK2 and GluK5 was unaltered in GluK2(Q) mice (Figure 4A) (GluK1: WT=100 ± 8.6%, GluK2(Q)=95.7. ± 8.5%, unpaired t-test, p=0.73; GluK2: WT=100 ± 8.1%, GluK2(Q)=140.6 ± 18.5 unpaired t-test, p=0.07; GluK5: WT=100.0 ± 9.8%, GluK2(Q)=108.4 ± 10.8%, unpaired t-test, p=0.57).

**Figure 4:**
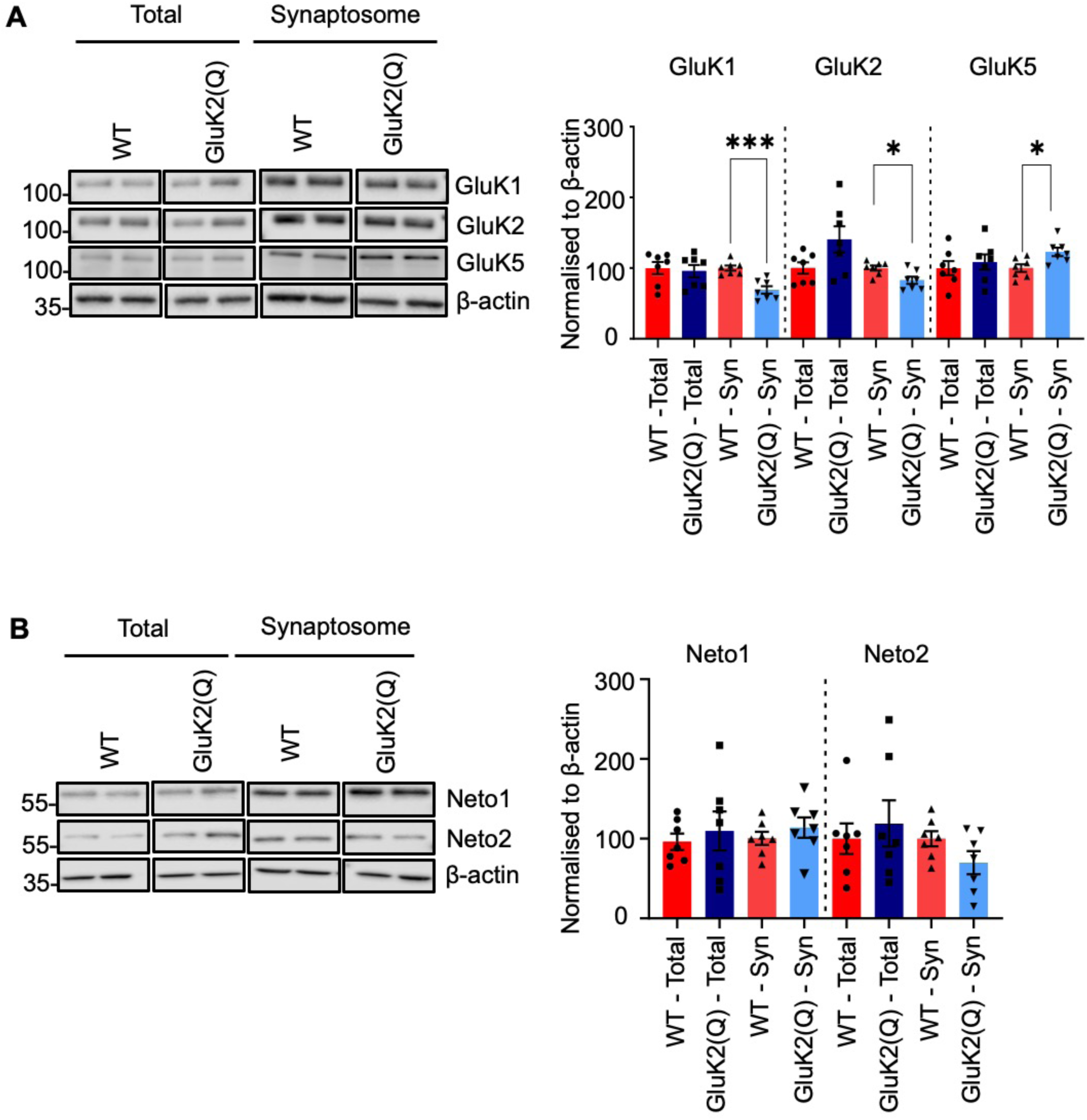
Altered synaptic KAR subunit expression in GluK2(Q) mice. **(A)** Representative Western blots of total and synaptosomal fraction samples from a single cerebral hemisphere of WT or GluK2(Q) mice for KAR subunits (left) . Quantification of proteins expressed as percentage of WT protein after normalizing to β -actin (right). β-actin was used as a loading control. N=7 animals; ns p>0.05, *p<0.05, **p<0.01; unpaired t-test with Welch’s correction. **(B)** Representative Western blots of total and synaptosomal fraction samples from a single cerebral hemisphere of WT or GluK2(Q) mice for auxiliary KAR subunits Neto1 and Neto2 (left) Quantification of proteins expressed as percentage of WT protein after normalizing to β-actin (right). β-actin was used as a loading control N=7; ns p>0.05, *p<0.05, **p<0.01; unpaired t-test with Welch’s correction.

In synaptosomes, however, the expression of GluK1 and GluK2 was reduced whereas expression of GluK5 was increased (Figure 4A) (GluK1: WT=100 ± 351%, GluK2(Q)=69.0 ± 5.3%, unpaired t-test, p=0.0006; GluK2: WT=100 ± 3.5%, GluK2(Q)=82.7 ± 5.1, unpaired t-test, p=0.002; GluK5: WT=100.0 ± 5.3%, GluK2(Q)=122 ± 6.0%, unpaired t-test, p=0.01). These data indicate that the enhanced pre- and postsynaptic ionotropic KAR function demonstrated in (Figures 1&2) are likely attributable to the increased conductance of GluK2(Q)-containing KARs, rather than an increase in the number of synaptic KARs.

Since most postsynaptic KARs comprise heteromeric combinations of GluK2/GluK5 and the auxiliary subunits Neto1 and/or Neto2 ^53^, we stripped the blots and reprobed for Neto1 and Neto2. Total (Figure 4B) (Neto1: WT=100 ± 10.5%, GluK2(Q)=114.0 ± 25.4%, unpaired t-test, p=0.62; Neto2: WT=100 ± 19.1%, GluK2(Q)=118.9 ± 29.0%, unpaired t-test, p=0.59) and synaptosomal expression of Neto1 and Neto2 were unchanged (Figure 4B) (Neto1: WT=100 ± 8.17%, GluK2(Q)=113.4 ± 12.6%, unpaired t-test, p=0.39; Neto2: WT=100 ± 9.4%, GluK2(Q)=69.8 ± 14.4%, unpaired t-test, p=0.11). We note, however, that because preparing MF-enriched synaptosomes would be extremely challenging these synaptosomes represent a heterogeneous population of synapses, and therefore changes in specific subsets of synapses might not be detected.

### Reduced synaptic AMPAR expression in GluK2(Q) mice

Given that GluK2(Q) mice display a reduction in the EPSC_AMPA_ (Figure 1D) we next stripped the same blots and reprobed for total and synaptic expression of AMPARs. Intriguingly, total and synaptosomal levels of the AMPAR subunits GluA1 and GluA3 were reduced (Total - GluA1: WT=100 ± 2.9%, GluK2(Q)=67.4 ± 8.7%, unpaired t-test, p=0.009; GluA3: WT=100 ± 6.5%, GluK2(Q)=37.3 ± 2.8%, unpaired t-test, p=0.006) (Synaptosomal - GluA1: WT=100 ± 4.8%, GluK2(Q)=65.1 ± 3.0%, unpaired t-test, p=0.0001; GluA3: WT=100 ± 4.2%, GluK2(Q)=52.7 ± 3.3%, unpaired t-test, p=0.002) (Figure 5A). In contrast, levels of GluA2 were unchanged in both total and synaptosome fractions (Total: WT=100 ± 9.9%, GluK2(Q)=73.4 ± 9.6%, unpaired t-test, p=0.08; synaptosome: WT=100 ± 7.7%, GluK2(Q)=118.2 ± 23.4%, unpaired t-test, p=0.4).

**Figure 5:**
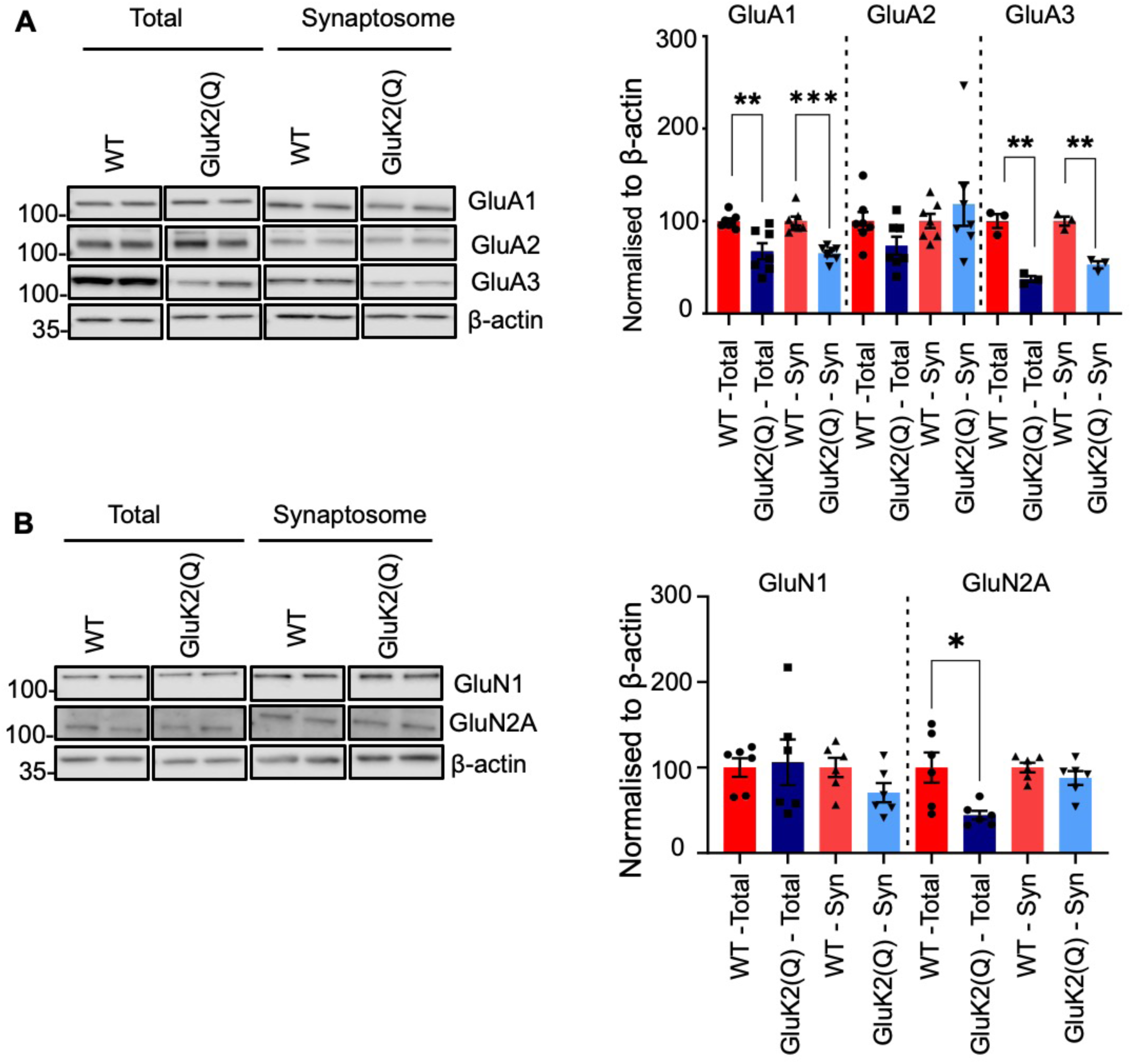
Reduced synaptic AMPAR expression in GluK2(Q) mice. **(A)** Representative Western blots of total and synaptosomal fraction samples from a single cerebral hemisphere of WT or GluK2(Q) mice for the AMPAR subunits GluA1-3 (left). Quantification of proteins expressed as percentage of WT protein after normalizing to β-actin (right). β-actin was used as a loading control. N=7 animals; ns p>0.05, *p<0.05, **p<0.01; unpaired t-test with Welch’s correction. **(B)** Representative Western blots of total and synaptosomal fraction samples from a single cerebral hemisphere of WT or GluK2(Q) mice for the NMDAR subunits GluN1 and GluN2A (left). Quantification of proteins expressed as percentage of WT protein after normalizing to β-actin (right). β-actin was used as a loading control N=6 animals (B); ns p>0.05, *p<0.05, **p<0.01; unpaired t-test with Welch’s correction.

We also tested NMDARs and found that although a reduction in the total levels of synaptically localised GluN2A was observed (Figure 5B) (GluN1: WT=100 ± 10.8%, GluK2(Q)=106.1 ± 26.7%, unpaired t-test, p=0.8; GluN2A: WT=100 ± 17.7%, GluK2(Q)=44.2 ± 5.2%, unpaired t-test, p=0.02), no changes were detected in the synaptic expression of GluN2A or the obligatory NMDAR subunit GluN1 ^54^ in GluK2(Q) mice (Figure 5B) (GluN1: WT=100 ± 11.2%, GluK2(Q)=70.6 ± 11.2%, unpaired t-test, p=0.09; GluN2A: WT=100 ± 5.5%, GluK2(Q)=87.7 ± 8.2%, unpaired t-test, p=0.2).

These data indicate that GluK2 Q/R editing selectively regulates the basal synaptic levels of both KARs and AMPARs, but not NMDARs.

### Impaired LTP at Schaffer collateral-CA1 synapses in GluK2(Q) mice

Since we observed a global reduction in the levels of AMPARs in the GluK2(Q) mice, we next characterised CA1 synapses by stimulation of a single presynaptic Schaffer-collateral axon at 1Hz in acutely prepared hippocampal slices ^46^. Postsynaptic responses were recorded from CA1 pyramidal neurons in whole-cell voltage clamp configuration in the presence of picrotoxin (50µM) and D-APV (50µM) to block GABA_A_Rs and NMDARs. As described for MF-CA3 synapses (Figure 1), stimulation intensity of presynaptic axons was increased until a postsynaptic response was observed (Figure 6A). The percentage of trials that failed to evoke a synaptic response (failure rate) was the same for both WT and GluK2(Q) mice (Figure 6B) (WT=0.60 ± 0.05%, GluK2(Q)=0.69 ± 0.04%; unpaired t-test, p=0.2). However, consistent with MF-CA3 synapses and the loss of AMPARs from the synaptosomes, the average AMPAR-EPSC amplitude for successful trials was reduced in GluK2(Q) mice (Figure 6C) (WT=28.8 ± 1.7pA, GluK2(Q)=19.2 ± 1.8pA; unpaired t-test, p=0.002).

**Figure 6:**
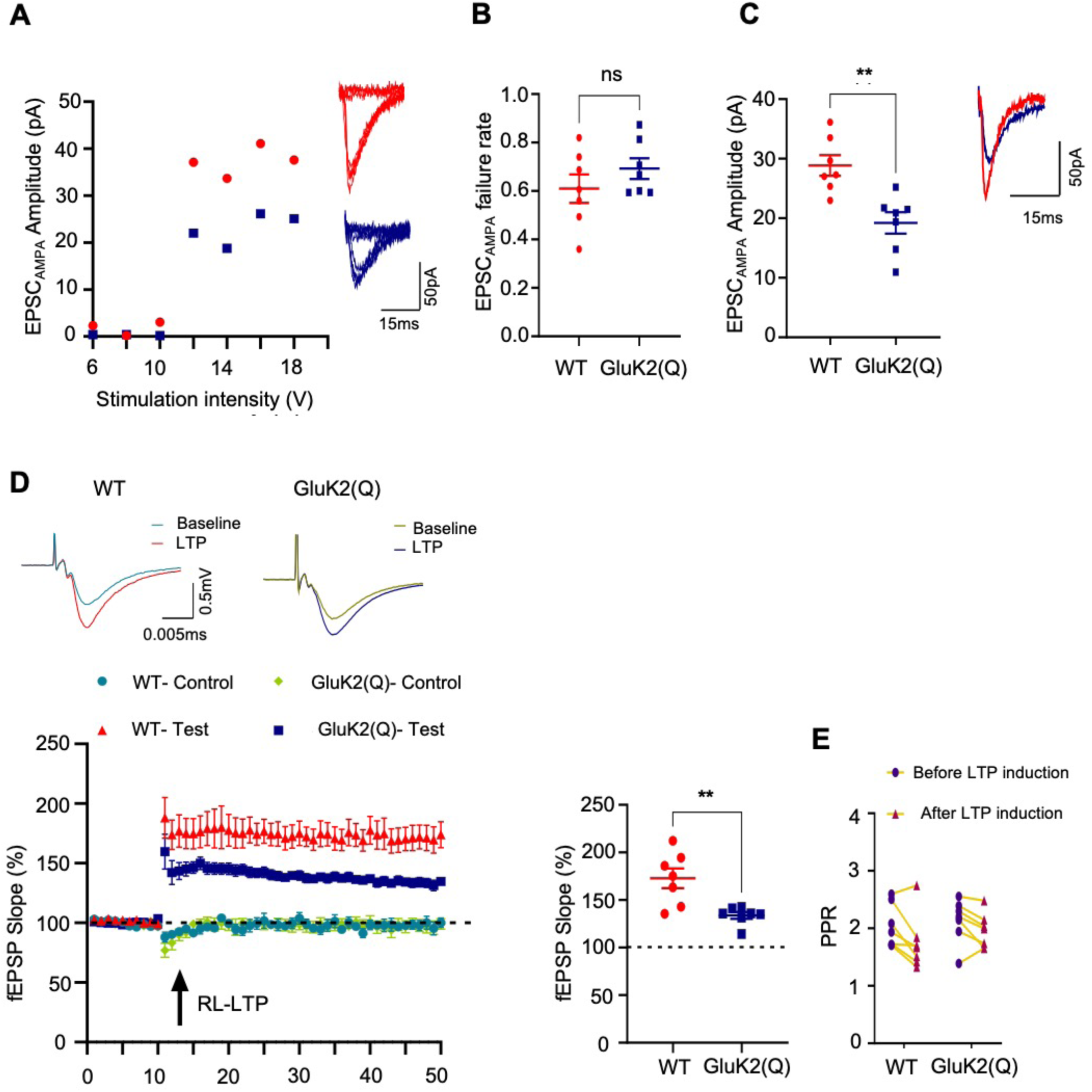
Impaired LTP at Schaffer collateral-CA1 synapses in GluK2(Q) mice. **(A)** Representative data from a minimal stimulation experiment in CA1 showing stimulation intensity threshold for evoking responses from excitation of a single axon fibre (left). Eight superimposed consecutive traces showing synaptic successes and failures for minimal stimulation for WT (top right) and GluK2(Q) mice (bottom right). **(B)** Quantification of probability of failures for AMPAR responses out of 150 stimulations in WT and GluK2(Q) mice. N=6, n=7 cells for WT and N=5, n=7 cells for GluK2(Q) mice; ns p>0.05; unpaired t-test with Welch’s correction. **(C)** Quantification of average amplitude of EPSC_AMPA_ (left). Representative trace showing EPSC_AMPA_ in WT and GluK2(Q) mice (right). N6, n=7 cells for WT and N=5, n=7 cells for GluK2(Q) mice; ns p>0.05, *p<0.05, **p<0.01; unpaired t-test with Welch’s correction **(D)** Representative traces showing field EPSPs (fEPSP) before and after LTP induction (21-30 min) in WT (top left) and GluK2(Q) mice (top right). Timeline showing fEPSP slope expressed as percentage of baseline subjected to Ripple-Like (RL)-LTP induction (arrow) (bottom left). Normalised fEPSP slope in test pathway 21-30 min after LTP induction in WT and GluK2(Q) mice (bottom right). . N=4 animals, n=7 cells; ns p>0.05, **p<0.002, ***p<0.0002, ****p<0.0001; Un-paired t-test with Welch’s correction **(E)** Paired-pulse ratio in WT and GluK2(Q) mice before and after LTP induction. N=4 animals, n=7 cells; ns p>0.05, **p<0.002, ***p<0.0002, ****p<0.0001; Two-way ANOVA with Sidak’s multiple comparison test.

These results further confirm the role of GluK2 Q/R editing in maintaining AMPAR-mediated synaptic transmission and indicate that this phenomenon occurs at different hippocampal synapses.

Since GluK2 editing prevents Ca^2+^ entry through the KAR ion channel we wondered if predominant expression of unedited GluK2(Q), which traffic more efficiently and gate Ca^2+^, would reveal EPSC_KAR_ at Schaffer collateral synapses. However, no EPSC_KAR_ were detected following bursts of 3 stimuli at 167Hz given to Schaffer collaterals to drive multiple axons (Supplementary Figure S2).

We next tested if the reduced synaptic expression of GluA1- and GluA3-containing AMPARs observed in GluK2(Q) mice impacted on the expression of long-term potentiation (LTP) at Schaffer collateral-CA1 synapses, a well characterized NMDAR-dependent form of LTP. Extracellular field potential recordings from acute hippocampal slices revealed that high frequency stimulation that replicates the *in vivo* patterns of hippocampal sharp-wave/ripple-like (RL) activity ^24,55^ induced robust LTP of AMPAR-mediated EPSPs in WT mice but LTP was significantly reduced in GluK2(Q) mice (Figure 6D) (WT: 172.8 ± 10.4% in test pathway vs 97.7 ± 3.8% in control pathway; unpaired t-test, p=0.0002; GluK2(Q): 133.8 ± 3.6% in test pathway vs 97.2 ± 4.9% in control pathway; unpaired t-test; p=0.0001; comparison between test pathways, unpaired t-test, p=0.0087). The paired-pulse ratio remained unchanged after induction of LTP in both WT and GluK2(Q) mice (Figure 6E) (WT=2.04 ± 0.14 baseline, 1.74 ± 0.17 after LTP, p=0.325; GluK2(Q)=2.13 ± 0.14 baseline, 1.95 ± 0.11 after LTP, p=0.65; Two-way ANOVA with Sidak’s multiple comparison test). These data suggest that preventing KAR Q/R editing reduces basal synaptic AMPAR expression and impairs the synaptic recruitment of AMPARs required for LTP.

## Discussion

GluK2 Q/R editing is developmentally ^37,56^ and activity-dependently regulated ^32,33^ to modulate and accommodate distinct synaptic and network diversity. However, how GluK2 editing impacts on KAR function and subsequent downstream AMPAR function has not been explored. To address these outstanding questions we compared WT mice, which contain <15% unedited GluK2(Q), and GluK2 Q/R editing deficient (GluK2(Q)) mice that have >95% unedited GluK2(Q) ^41^.

In recombinant expression systems KARs containing unedited GluK2(Q) have a much greater single channel conductance than those containing edited GluK2(R) (∼150ps compared to <10ps) ^40^. We therefore predicted that postsynaptic ionotropic KAR function would be enhanced in GluK2(Q) mice. We found this to be the case, but the increase in EPSC_KA_ we observed (WT=4.66 ± 0.63pA, GluK2(Q)=7.31 ± 0.50pA) was markedly less than expected based on the data from heterologous expression systems. However, it should be noted that heterologous expression systems can lack neuron specific proteins (e.g., Netos, KRIP6) that are known to affect the gating of the receptors. In addition, we are unaware of any information on how heteromeric KARs containing GluK2(Q) or GluK2(R) subunits behave when combined with GluK4/GluK5.

Similarly, we anticipated that the increased conductance and Ca^2+^ permeability of GluK2(Q)-containing KARs should boost presynaptic KAR function, resulting in enhanced short-term facilitation at both 50ms (PPF) and 1s (FF) timescales. This is indeed what we found, but we also observed an increase in failure rate in the minimal stimulation experiments. These data suggest an additional factor of reduced basal probability of release, or number of release sites, within the large presynaptic mossy fibre boutons, which on its own is predicted to increase presynaptic facilitation. Minimal stimulation at 3Hz is sufficient to engage KAR-mediated FF and would therefore be expected to produce a lower failure rate in GluK2(Q) mice. We observed the opposite effect, suggesting that the reduction in basal probability of release is substantial. Our data cannot distinguish between reduced probability of release and reduced number of release sites so further anatomical investigation would be necessary to address this. Nonetheless, overall, our data indicate that GluK2(Q) mice exhibit enhanced pre- and postsynaptic KAR function.

In stark contrast to their enhanced ionotropic KAR function, GluK2(Q) mice show reduced metabotropic function measured by inhibition of I_sAHP_ at MF-CA3 synapses ^15,26,27,52^. Possible explanations for the diminished metabotropic KAR signalling in GluK2(Q) mice include a reduction in expression of GluK1 and GluK2 KAR subunits and/or Q/R editing state-dependent conformational changes which regulate metabotropic signaling. It remains unclear and controversial which, and how, specific KAR subunits contribute to KAR metabotropic signalling. Indeed, it has been proposed by different groups that GluK1, GluK2, or GluK5 are required for G protein coupling and metabotropic effects ^15,21,24,31,57^. Thus, although we cannot draw definitive mechanistic conclusions, our data demonstrate that GluK2(Q) mice show reduced metabotropic KAR signalling and, at the same time, enhanced ionotropic function.

A critical role for KAR signaling is activity-dependent regulation of both KAR and AMPAR surface expression. For example, depending on the extent of activation, KARs can enhance or reduce AMPAR surface expression to evoke LTP or LTD via metabotropic or ionotropic signalling, respectively ^24,35^. Our data also show that GluK2 editing impacts on NMDAR-induced LTP at Schaffer collateral synapses in CA1, with GluK2(Q) mice exhibiting reduced LTP. GluK2(Q) mice also had reduced AMPAR-EPSCs at mossy fibre and Schaffer collateral synapses and lower GluA1 and GluA3 levels in synaptosomal fractions, suggesting that KARs not only regulate activity-dependent AMPAR trafficking but may also maintain basal synaptic levels of Ca^2+^-permeable AMPARs. It should be noted that activity-dependent regulation of KAR alters the synaptic expression of GluA1 and GluA2 AMPARs, while GluK2 editing specifically alters the basal synaptic expression of GluA1 and GluA3 containing AMPARs ^24,35^. Importantly, these changes were specific to AMPARs, since synaptic levels of the NMDAR subunits GluN1 and GluN2A were unchanged, indicating the loss of AMPARs is not due to wholesale changes in synaptic composition in the editing-deficient mice.

These findings demonstrate that KAR activity mediates the ‘tone’ of synaptic AMPARs at MF-CA3 and Schaffer collateral synapses and suggests a wider role where KAR signalling may set the tone for synaptic AMPAR composition more broadly. We speculate that this may be a homeostatic mechanism in which the presence of high-conductance, Ca^2+^-permeable GluK2(Q)-containing KARs causes a compensatory decrease in Ca^2+^-permeable GluA1/GluA3-containing AMPARs to balance synaptic responsiveness. This is similar to the observed effects of KAR signalling on expression of AMPARs during the development of synaptic circuits ^47,58^ and suggests that this developmental regulation extends into adulthood.

In healthy adult brain GluK2-containing KARs predominantly comprise edited GluK2(R) ^59^. Using mice that almost exclusively express only GluK2(Q) we show that the ionotropic/metabotropic balance of KAR signalling is radically altered by a lack of GluK2 editing. Based on these results, we propose that unedited GluK2(Q)-containing KARs primarily or exclusively function as ion channels with enhanced conductance for both mono and/or di-valent cations, whereas the edited GluK2(R)-containing KARs act as metabotropic receptors to regulate and maintain network activity.

These findings are important because GluK2 Q/R editing is subject to both developmental and activity-dependent control ^33^. Moreover, it has been reported that the proportion of edited GluK2(R) is increased to 85% in patients with a pharmaco-resistant temporal lobe epilepsy (TLE) ^59^, raising the possibility that increased inhibition of I_sAHP_, and thereby hyperexcitability, could underpin seizure generation. Taken together our data indicate that physiologically and pathologically relevant alterations in GluK2 editing may dynamically regulate KAR function, signalling mode, maintenance of functional neuronal networks, and set the threshold for the induction of plasticity. Thus, in conclusion, our results highlight that GluK2 Q/R editing acts as a previously unsuspected molecular switch that regulates the enigmatic dual-mode capability of KARs to operate via either ionotropic or metabotropic signalling, to initiate distinct and diverse downstream pathways.

## Acknowledgments

This work was supported by the BBSRC (BB/R00787X/1), the Leverhulme Trust (RPG-2019-191) and the Wellcome Trust (105384/Z/14/A). We are grateful to Prof. Susumu Tomita (Yale, USA) for the kind gift of NETO antibody.

## Author contributions

Conceptualization, JDN, KAW, JM, JMH.

Methodology, JDN, KAW, JM, CM.

Investigation, JDN, BPY, KAW.

Writing JDN, KAW, JM, CM, JMH.

Original Draft, and Writing, JDN, KAW, JM, JMH.

Review & Editing, All authors.

Funding Acquisition, JMH.

Resources, BV.

Supervision, JMH, KAW, JRM;

## Supplemental information

**Supplementary Figure S1:**
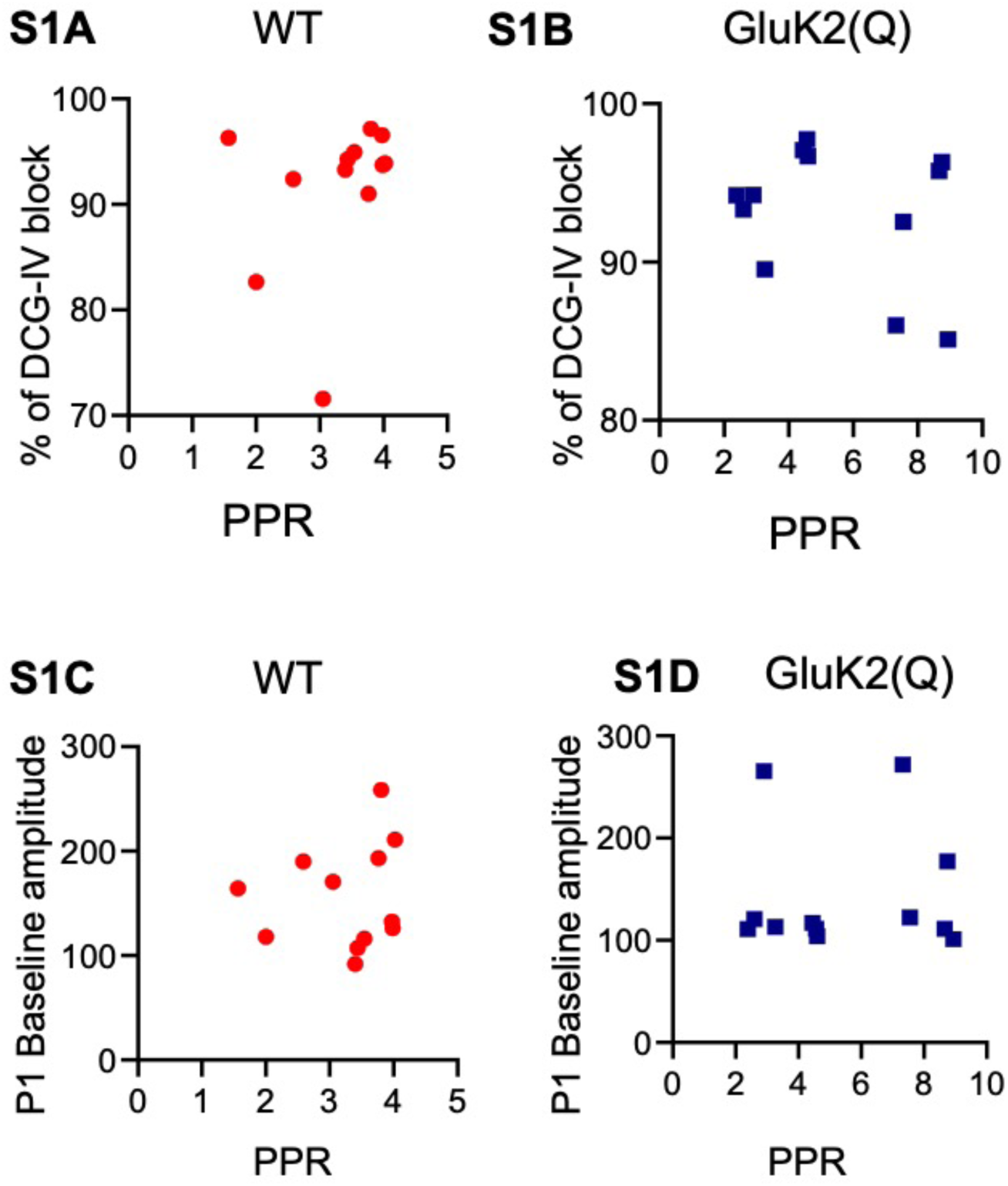
Pearson’s correlation between PPR and percentage of DCG-IV block in WT **(S1A) and** GluK2(Q) mice **(S1B)**. PPR and P1 baseline amplitudes of WT **(S1C)** and GluK2(Q) mice **(S1D).** WT, N=7, n=13 cells; Tg, N=8, n=15 cells.

**Supplementary Figure S2:**
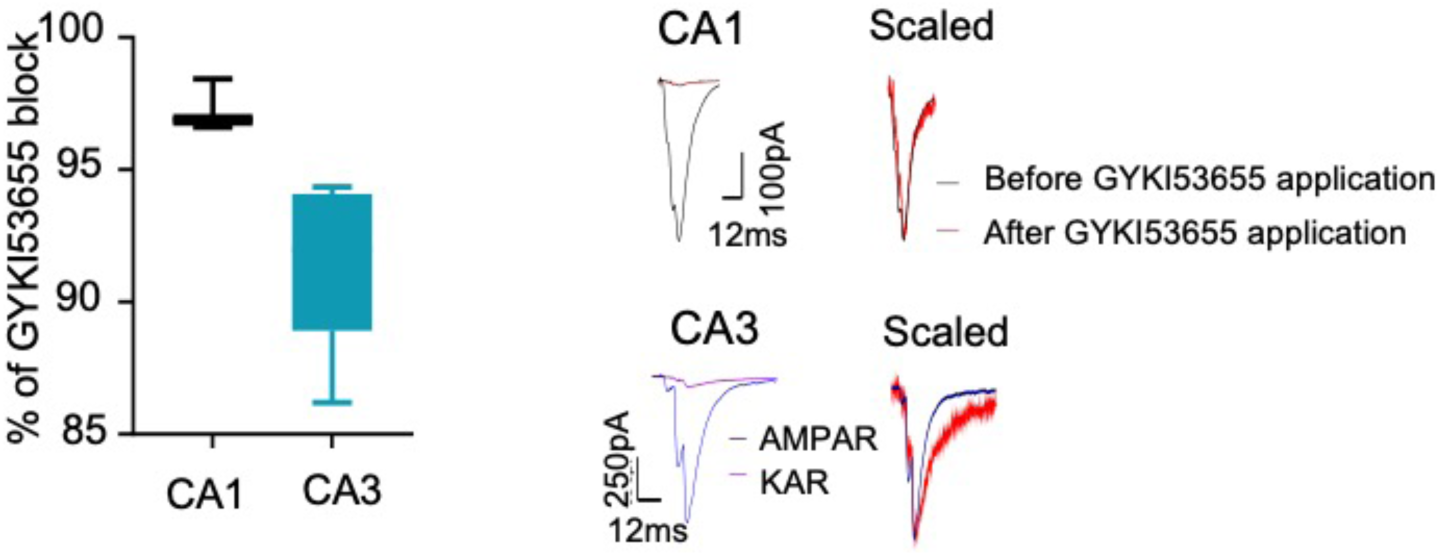
Quantification of percentage of GYKI53655 block in CA1 and CA3 synapses in GluK2(Q) mice **(left)**. Representative traces showing postsynaptic EPSCs elicited by bursts of stimuli in CA1 **(top middle)** and CA3 **(bottom middle)** of GluK2(Q) mice before and after 10 mins of GYKI5655 (40µM) application to block AMPA receptors. A larger percentage of the response remains after GYKI53655 in CA3. Scaled response before and after GYKI53655 application in CA1 **(top right)** and CA3 **(bottom right)**. The remaining CA1 currents show identical decay kinetics indicating a residual AMPAR response, while CA3 currents post-GYKI53655 application show a slow decay kinetics consistent with KAR responses.

## Methods

### Animals

GluK2 editing-deficient ECS mice and their WT counterparts (129Sv strain) were created at the Salk Institute by mutating the intronic editing complementary sequence (ECS) in the *grik2* gene that directs ADAR2-mediated codon substitution in the GluK2 pre-mRNA. ^41^.

The mice were housed in groups of 2-4 in standard individually ventilated (IVC) cages in rooms with temperature maintained between 19-23°C and with 12h light and dark cycles. Cages had sawdust, paper nesting and an enriched environment (wooden chews, cardboard tubes etc.). Pups of age P14-P21 were used for the experiments irrespective of sex.

All the animal experiments and procedures were performed in compliance with the UK Animal Scientific Procedures act (1986) and were guided by the Home Office Licensing Team at the University of Bristol. All animal procedures relating to this study were approved by the Animal Welfare and Ethics Review Board at the University of Bristol (approval number UIN/18/004).

All experiments and analysis were performed blinded to mouse genotype.

### Acute hippocampal slice preparation

Cervical dislocation followed by decapitation were performed on P14-21 male and female WT and GluK2(Q) mouse pups. The brain was removed and placed in ice-cold sucrose slicing solution (in mM: Sucrose, 205; KCl, 2.5; NaHCO_3_, 26; NaH_2_PO_4_, 1.25; D-Glucose, 10; CaCl_2_, 0.5; MgCl_2_, 5) saturated with 95% O_2_ and 5% CO_2_. Hippocampi were carefully removed and transverse sections of 400µm thickness for whole-cell recordings and 500 µm for field recordings were obtained using a vibratome (Leica VT 1200s). Slices were kept for recovery in a slice holder containing artificial cerebrospinal fluid (aCSF; in mM: NaCl, 124; KCl, 3; NaHCO_3_, 24; NaH_2_PO_4_, 1.25; D-Glucose, 10; MgSO_4_, 4; CaCl_2_, 4) for whole-cell recordings and (aCSF; in mM: NaCl, 124; KCl, 3; NaHCO_3_, 24; NaH_2_PO_4_, 1.25; D-Glucose, 10; MgSO_4_, 2; CaCl_2_, 2) for field recordings saturated with 95% O_2_ and 5% CO_2_ at 37°C for 20 mins and later transferred to room temperature for at least 30 mins before performing experiments.

### Electrophysiology recordings

#### Whole cell recordings

Hippocampal slices were placed in a submerged holding chamber continuously perfused with oxygenated aCSF at 36.5°C at a flow rate of 3ml per minute. Hippocampal CA3 pyramidal cells were visually identified using DIC optics and patch-clamped in whole-cell configuration using a pulled Harvard borosilicate glass capillary of resistance 5-7M MΩ filled with either caesium-based whole-cell solution (in mM: NaCl, 8; CsMeSO_4_, 130; HEPES, 10; EGTA, 0.5; MgATP, 4; NaGTP, 0.3; QX314.Cl, 5; Spermine, 0.1) or K-Gluconate based (in mM: NaCl, 8; KGluconate, 135; HEPES, 10; EGTA, 0.2; MgATP, 2; NaGTP, 0.3) for I_sAHP_ experiments.

The cells were held in voltage clamp mode and evoked EPSCs were obtained by stimulating the mossy fibre pathway with a bipolar stimulating electrode placed in the dentate gyrus hilus layer (or glass monopolar electrode for minimal stimulation experiments). Picrotoxin (50µM) (Sigma: P1675) was included in the aCSF to inhibit GABA_A_ receptors (except for CA1 field recordings). Cells with series resistance above 30 MΩ or where series resistance changed by >20% were excluded from analysis. To confirm the purity of mossy fibre inputs, the group-II mGluR agonist DCG-IV (2µM) (Tocris: 0975/1) was bath applied for 5-10 mins at the end of experiments with mossy fibre stimulation ^27^. Recordings were only included in analysis if DCG-IV reduced EPSCs by >70%.

Data were digitized at 10kHz, and low-pass filtered at 2kHz using CED Micro 1401-4 A-D acquisition unit and Axon patch 200B amplifier (Molecular devices). All recordings were obtained using CED Signal 5 software.

CED Signal acquisition software was used to analyze the recorded data. Mean responses were obtained every minute by averaging consecutive traces. EPSC amplitudes were measured from the averaged traces and normalized to the mean EPSC amplitude of baseline.

#### Minimal stimulation

##### CA3 pyramidal cells

Cells were voltage clamped at -60mV and MF-EPSCs were evoked by moving a mono-polar stimulating electrode filled with caesium-based whole-cell solution (in mM: NaCl, 8; CsMeSO_4_, 130; HEPES, 10; EGTA, 0.5; MgATP, 4; NaGTP, 0.3; QX314.Cl, 5; Spermine, 0.1) around the inner border of dentate gyrus granule cells until a response was observed. Stimulation intensity was adjusted just above the threshold for activation of a synaptic response. Consecutive traces were recorded at a frequency of 3Hz. No prominent polysynaptic activation was observed using this low intensity stimulation. AMPAR-EPSCs were measured for 5-10 mins in the presence of D-APV (50µM) for a minimum of 150 trials. Subsequently, GYKI53655 (40μM) was applied to block AMPAR responses and KAR-EPSCs were measured after 15 mins of GYKI53655 application and for 5-10 mins and a minimum of 150 trials. !_decay_ for AMPAR and KAR-EPSCs were calculated by single exponential curve fitting feature in CED signal software.

##### CA1 pyramidal cells

Cells were voltage clamped at -60mV and Schaffer collaterals were evoked by moving a mono-polar stimulating electrode filled with caesium-based whole-cell solution (in mM: NaCl, 8; CsMeSO_4_, 130; HEPES, 10; EGTA, 0.5; MgATP, 4; NaGTP, 0.3; QX314.Cl, 5; Spermine, 0.1) around the stratum radiatum until a response was observed. Stimulation intensity was adjusted just above the threshold for activation of a synaptic response. Consecutive traces were recorded at a frequency of 1Hz. No prominent polysynaptic activation was observed using this low intensity stimulation. AMPAR-EPSCs were measured for 5-10 mins in the presence of D-APV (50µM) for a minimum of 150 trials.

#### KAR/AMPAR ratio

CA3 pyramidal neurons were voltage clamped at -60mV in the presence of D-APV (50µM) (Hello Bio: HB0225). Mossy fibres were stimulated with a burst of 3 stimuli at 167Hz every 20s to evoke AMPAR/KAR-EPSCs. Stable AMPAR-EPSCs were recorded for 20 minutes and then KAR-EPSCs were recorded for 20 mins in the presence of the AMPAR antagonist GYKI53655 (40μM) (Hello Bio: HB0312). Amplitude of AMPAR-KAR peaks and KAR peaks were measured individually and the ratio of KAR-EPSC to AMPAR-EPSC were calculated.

#### Paired-pulse and frequency facilitation

CA3 neurons in acute hippocampal slices were voltage clamped at -70mV in the presence of picrotoxin (50µM). To measure PPF, EPSCs were evoked by pairs of stimuli to mossy fibres at an inter-stimulus interval of 50ms, every 20s. Paired-pulse ratios were obtained by averaging amplitudes of P1 peak to P2 peak. For FF experiments, single stimuli were given at 0.05Hz for 10 mins before stimulation frequency was increased to 1Hz for 1 min. FF ratios were obtained by averaging the last 20 frames of P1 amplitude at 0.05Hz with the middle 40 frames at 1Hz stimulation. After this, stimulation frequency was returned to 0.05Hz. DCG-IV (2μM) was then applied for at least 5 mins to assess purity of MF input.

#### Slow afterhyperpolarizations (I_sAHP_)

For I_sAHP_ recordings, CA3 pyramidal neurons were voltage clamped at -50mV in the presence of picrotoxin (50µM), D-APV (50µM), CGP55845 (1µM) (Hello Bio: HB0960) and GYKI53655 (40μM) . Glass electrodes were filled with whole-cell solution (in mM: NaCl, 8; KGluconate, 135; HEPES, 10; EGTA, 0.2; MgATP, 2; NaGTP, 0.3). I_sAHP_ were induced every 20s by applying a depolarising voltage step to 0mV for 200ms and I_sAHP_ amplitude was measured 300ms after returning the membrane potential to -50mV to avoid measurement of medium afterhyperpolarization (I_mAHP_). Synaptic activation of KARs was induced by bursts of 10 stimuli at 25Hz to the mossy fibres 500ms prior to the induction of I_sAHP_.

#### Field potential recordings

Extracellular field potentials (fEPSPs) were recorded from stratum radiatum in CA1 using a 3-5 MΩ glass pipette filled with aCSF. Two stimulation electrodes (bipolar) were positioned on opposite sides of the recording electrodes equidistant from the pyramidal layer to evoke two independent inputs (Stim1 and Stim2). LTP induction protocol was delivered only to Stim1 and was alternately positioned closer to the CA3 region or to subiculum in different recordings. Paired stimuli (50ms inter-stimulus interval) were given every 10s to each pathway, alternating between the control and test pathway (Stim1 and Stim2). LTP consisted of 20 bursts of 20 stimuli at 200Hz given every 5s. The recordings were performed in the absence of picrotoxin. fEPSP slopes were measured using CED signal software. The slopes are displayed as a percentage of 10 min baseline. For LTP quantification the values were obtained 21-30 min after LTP induction.

### Synaptosomal preparations and Western blotting

Synaptosomes were prepared from cerebral hemisphere of P14-21 pups using Syn-PER^TM^ Reagent (ThermoFisher Scientific: 87793). After cervical dislocation followed by decapitation of the pups, the brain was removed and cut into two halves (along the cerebral hemispheres). The cerebellum was discarded. Each cerebral hemisphere from the pup was weighed and transferred to a glass homogenizer and the required amount of Syn-PER reagent was added to the tissue (10ml of reagent per gram of tissue). The tissue was homogenized on ice with slow stokes (∼10 strokes). The homogenate was transferred to a fresh centrifuge tube and centrifuged at 1200 x g for 10 min at 4°C. The supernatant was transferred to a fresh tube and the pellet was discarded. The supernatant was centrifuged again at 15,000 x g for 20 min at 4°C. The supernatant was discarded, and the pellet was resuspended in 1-2ml of Syn-PER reagent. To the synaptosomal fraction, Triton x-100 and SDS were added to a final concentration of 1% and 0.1%, respectively, and left at 4°C on a rotating wheel for 1hr to lyse synaptosomes and solubilize membrane proteins. The samples were then centrifuged at 16,000 x g for 20 mins to remove insoluble material. Protein quantification was performed on the samples and the final samples for Western blotting were prepared by adding 2X sample buffer and boiling at 95°C for 10 mins.

The samples prepared were separated based on molecular weight using Sodium dodecyl sulphate-poly acrylamide gel electrophoresis (SDS-PAGE). The gel comprised a 10% acrylamide resolving gel (375mM Tris-HCL pH 8.8, 10% acrylamide, 0.1% SDS, 0.1% APS and 0.01% TEMED) and 5% stacking gel (125mM Tris-HCL pH 6.8, 5% acrylamide, 0.1% SDS, 0.1% APS, 0.01% TEMED) .The gels were transferred to PVDF membrane and blocked in 5% skimmed milk in PBS-T (0.137M NaCl, 2.7mM KCl, 10mM Na_2_HPO_4_, 2mM K_2_HPO_4_, pH to 7.4 with HCl, 0.001% Tween-20) for 1 hr at RT and blotted overnight at 4°C in the same blocking solution with the following antibodies: Rabbit Anti-GluK1 (1:1000; Millipore: 07-258), Rabbit Anti-GluK2/3 (1:1000; Millipore: 04-921), Rabbit Anti-GluK5 (1:1000; Millipore: 06-315), in-house Rabbit Anti-Neto1 (1:1000; a kind gift from Prof. Susumu Tomita (Yale, USA)), Rabbit Anti-Neto2 (1:1000; Abcam: Ab109288), Rabbit Anti-GluA1 (1:1000; Millipore: AB1504), Mouse Anti-GluA2 (1:1000, BD Pharmighen: 556341), Rabbit Anti-GluA3 (1:1000; Alomone labs: AGC-010), Rabbit Anti-NMDAR1 (1:1000; Abcam: Ab109182), Rabbit Anti NR2A (1:1000; Abcam: Ab124913), Mouse Anti-β-Actin (1:10000; Sigma: A5441). The HRP-Conjugated secondary antibodies((Anti-Rabbit (raised in goat) and anti-Mouse (raised in goat)) were from Merck. . The antibodies were used at 1:10,000 dilution in 5% milk in PBS-T.

For each experiment, the signal for each condition was divided by the signal from the loading control for that experiment (β-actin). This analysis was performed for each replicate experiment, and for presentation purposes, the mean of the control condition set to 100%.

### Data analysis

Data are plotted as mean ± SEM. ‘N’ - Number of animals used, ‘n’ - number of cells. ANOVA, or paired or un-paired student t-test were used for statistical analysis and stated in the figure legends with respective p values. All statistical tests were performed using GraphPad Prism version 9.3.

